# Client-scaffold interactions suppress aggregation of a client protein in model condensates

**DOI:** 10.1101/2025.04.11.648422

**Authors:** Rashik Ahmed, Rhea P. Hudson, Julie D. Forman-Kay, Lewis E. Kay

## Abstract

Many studies have shown that sequestration of client proteins into condensates locally increases their concentrations and/or modulates their conformational landscapes to promote aberrant aggregation. Far fewer examples have emerged where the proteinaceous condensed phase environment protects clients from aggregation. Here, we show that a condensate scaffolded by the C-terminal disordered region of Cell Cycle Associated Protein 1 (CAPRIN1) suppresses aggregation of the Fused in Sarcoma (FUS) RNA Recognition Motif (RRM) client, both components of stress granules. Although FUS RRM aggregation is mediated through the unfolded ensemble, comparative NMR studies of the FUS RRM outside and within the condensate establish that CAPRIN1 condensates attenuate FUS RRM aggregation despite locally increasing its concentration by 2-fold and significantly unfolding the domain. Regions of transient intermolecular contacts between unfolded FUS RRM protomers driving aggregation have been identified, including the hydrophobic segments spanning I287-I308 and G335-A369. Intermolecular NOE experiments recorded on the FUS RRM:CAPRIN1 condensate indicate that CAPRIN1 interacts with much of the unfolded FUS RRM, with regions of stronger contacts including the RRM sequences ^287^IFVQ^290, 296^VTIES^300, 322^INLY^325^, and ^351^IDWFDG^356^. These interactions collectively outcompete the homotypic contacts between unfolded FUS RRM clients driving aggregation. Our results demonstrate that condensate scaffold molecules can, in some cases, shield client interprotomer interactions, delaying or completely suppressing their aggregation.

**SIGNIFICANCE STATEMENT:** Numerous studies demonstrate that condensates can promote protein aggregation within cells, potentially leading to disease. Here we show that in some cases condensates protect against aggregation of client proteins within them, using a model system consisting of a pair of proteins that are found in stress granules. Protection occurs even though unfolded client polypeptide chains, normally associated with aggregation, increase significantly in concentration in the proteinaceous condensed-phase environment. Using solution NMR spectroscopy, we provide an atomic resolution map of the interactions between the client, an RNA recognition module from the protein FUS, and a phase-separating scaffold protein, CAPRIN1, that protect against aggregation. These findings broaden our understanding of the mechanisms by which condensates regulate cellular protein homeostasis.

## INTRODUCTION

The cellular milieu is, in part, organized into numerous non-membrane-bound compartments that form and dissipate in response to various cellular cues(1, 2). These compartments, referred to as biomolecular condensates, often emerge through the process of phase separation(1, 3). The high local concentration of biomolecules in condensates generates unique solvent environments that selectively enrich a subset of molecules. The biomolecules that comprise the condensate can be simplified into two classes: “scaffolds” which are necessary for the structural integrity of the condensate and “clients” that partition into the condensate but are not essential for their formation(1). Condensate composition can vary dramatically under different cellular conditions and change rapidly in response to signaling(4). Through such compositional control, condensates can regulate the specificity and kinetics of biochemical processes(5–7).

While it is clear that condensates play crucial roles in normal cellular function(8), growing evidence suggests that they may also contribute to disease pathogenesis. Indeed, condensation often leads to an increase in the local concentration of client proteins, including those that are aggregation-prone, which can accelerate the nucleation and growth of pathological protein aggregates. This has been observed for Fused in Sarcoma (FUS)(9), TDP-43(10), α-synuclein(11), heterogeneous nuclear ribonucleoprotein A1 (HNRNPA1)(12) and Tau(13) proteins linked to various neurodegenerative disorders. The unique solvent environment of condensates can also shift the conformational landscape of proteins towards states that are susceptible to aggregation, as we have recently demonstrated for the superoxide dismutase 1 (SOD1) client protein in condensates scaffolded by cell cycle associated protein 1 (CAPRIN1)(14) or others have established through phase separation of SOD1 itself(15). Therefore, it appears that condensates can enable access to previously inaccessible protein states that function as unstable intermediates on-pathway to formation of aberrant protein aggregates.

It is also possible to envision cases where condensate scaffold proteins stabilize protein client conformations such that they become less susceptible to aggregation, or where scaffolds directly compete for intermolecular client interactions that would otherwise promote aggregation. Indeed, several reports indicate that condensates can suppress or significantly slow down the aggregation of client proteins despite enhancing their local concentrations(16, 17). Notably, this behaviour is dependent on the scaffold proteins that assemble the condensate, with some scaffolds promoting client aggregation and others slowing it down, presumably through stabilization of the monomeric client form(17). However, it remains to be understood, at the atomic level, what the network of interactions involved in the solvation of client proteins inside condensates are and how such interactions can sometimes prevent aggregation.

Solution NMR spectroscopy is a powerful technique for studies of biomolecules within the highly dynamic environments of phase-separated condensates. Through appropriate labeling of the biomolecules of interest with NMR-active spins(18), coupled with sensitive experiments that are optimized for studies of high molecular weight particles(19– 21), it becomes possible to quantify – at atomic resolution – homotypic interactions between proximal client molecules or between scaffold proteins(22), as well as heterotypic client-scaffold interactions(14). Here, we use solution NMR to examine the conformational landscape of an aggregation-prone client protein inside a model scaffold condensate. Our system consists of the RNA recognition motif (RRM) domain of the stress granule protein FUS (Fig. 1A, *left*). The RRM domain dramatically enhances the aggregation of full-length FUS(23) and in isolation it spontaneously self-assembles into amyloid fibrils(24). The FUS RRM client was incorporated into condensates formed by the C-terminal low complexity region (S607 - Q707) of the stress granule protein CAPRIN1 (Fig.1A, *right*), referred to as CAPRIN1 in what follows. While the FUS RRM aggregates in buffer solution or in the dilute phase of the phase-separated FUS RRM:CAPRIN1 system, high concentrations of CAPRIN1, such as those found in the condensed phase, suppress aggregation. Remarkably, suppression of aggregation occurs despite a shift of the FUS RRM conformational landscape in the condensed phase towards unfolded states that normally are prone to aggregation. We identified numerous sites of intermolecular contacts between unfolded FUS RRM protomers using NMR experiments quantifying intensities of cross peaks in spectra performed as a function of FUS concentration. Notably, sites of heterotypic FUS RRM:CAPRIN1 interactions identified by recording NOESY spectra in the condensed phase significantly overlap with regions of homotypic FUS RRM-FUS RRM contacts. Collectively, our findings provide atomic-level insights into potential protective mechanisms of condensates for aggregation-prone proteins, whereby client interprotomer interactions that drive aggregation are effectively outcompeted by heterotypic interactions involving scaffold molecules and/or where contacts with scaffold proteins result in stabilizing client conformations that are unable to evolve into pathological aggregates.

**Figure 1.**
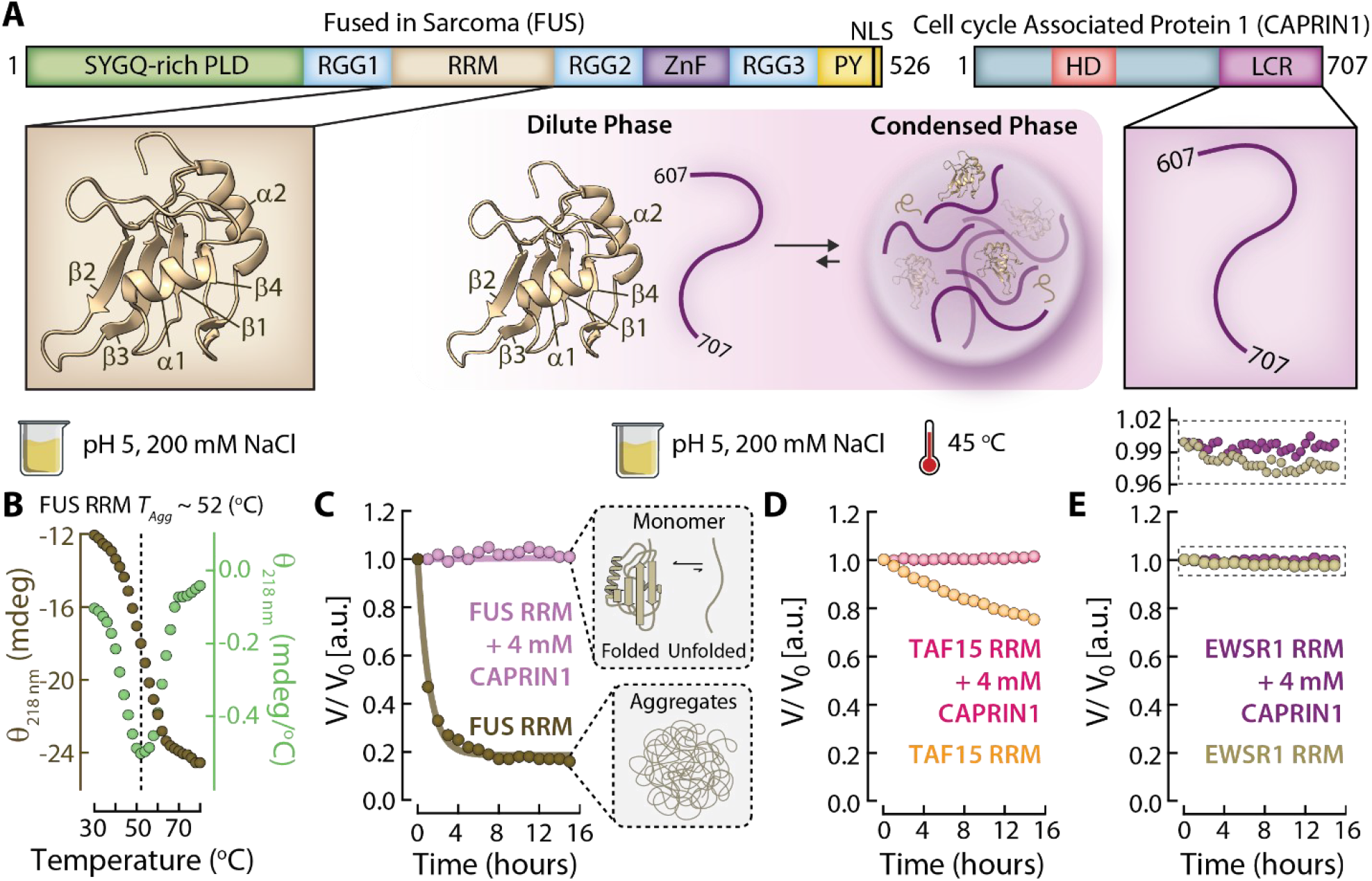
Mixed CAPRIN1: RRM solutions suppress aggregation of FUS, EWSR1, and TAF-15 RRMs. (**A**) Domain architecture of FUS (*left; PDB ID: 2LCW*) and CAPRIN1 (*right*). (**B**) Circular dichroism thermal melt monitoring the β-sheet signature at 218 nm of FUS RRM in MES pH 5 buffer with 200 mM NaCl (*brown*). Note that the low pH and high salt conditions used here drive aggregation of FUS RRM. The derivative of the thermal melt profile is in green, from which T_Agg_ ≈ 52 °C is measured. (**C**) Aggregation kinetics of 0.5 mM FUS RRM in the absence (*brown*) and presence of 4 mM CAPRIN1 (*purple*), as monitored by the loss of NMR-visible FUS RRM amide signals in 1D ^15^N-edited ^1^H NMR spectra. Spectra were recorded at 45 °C, 600 MHz. (**D-E**) As (**C**), except for the TAF-15 (**D**) and EWSR1 (**E**) RRMs, with color-coding as per the panel labels. The inset above **E** shows an enlarged view of the EWSR1 RRM kinetic profiles.

## RESULTS

### CAPRIN1 suppresses the aggregation of RRMs from the FUS, EWSR1, TAF15 (FET) family of proteins

We evaluated the thermal stability of FUS RRM under acidic conditions typically associated with cellular stress(25) using circular dichroism (CD) spectroscopy. The FUS RRM CD signal at 218 nm becomes increasingly negative with temperature indicating a conversion of the monomeric protein into beta-sheet rich aggregates with an approximate temperature for half-maximal aggregation (*T*_*agg*_) of 52 °C (Fig. 1B). We next monitored the aggregation of FUS RRM at temperatures associated with moderate heat stress *i*.*e*. 45 °C(26), yet below *T*_*agg*_, that allows for sufficient sampling of the aggregation profile in relatively short periods of time (< 1 day). Aggregation was probed by measuring the loss of NMR-visible FUS RRM amide signals in 1D ^15^N-edited ^1^H NMR spectra as a function of time using a sample that was initially at a concentration of 0.5 mM. Under these conditions, ∼80% of NMR visible FUS RRM is converted into NMR-invisible aggregates within ∼6 hours (Fig. 1C, *brown*). In contrast, addition of high concentrations of CAPRIN1 (4 mM) to 0.5 mM FUS RRM completely suppresses its aggregation (Fig. 1C, *purple*). This is also the case for TAF15 (Fig. 1D) and EWSR1 (Fig. 1E) RRMs, albeit with less conversion to aggregates compared to FUS. Notably, the significant decrease in aggregation kinetics of EWSR1 relative to the other FET RRMs, as quantified in our NMR experiments, is consistent with the higher *T*_*agg*_ value of 64 °C (an increase of ∼ 10 °C) measured for this domain (*SI Appendix*, Fig. S1).

### Intermolecular FUS RRM:FUS RRM interactions

Having established that CAPRIN1 is protective against FET RRM aggregation under acidic conditions involving moderate heat stress (Fig. 1C-E) we next sought to characterize, on a per-residue level, the interactions between FUS RRM molecules that contribute to aggregation. To this end, we prepared uniformly ^15^N and ^13^C-labeled FUS RRM samples at low (0.15 mM) and high (0.6 mM) concentrations in a pH 6 buffer for NMR experiments recorded at 45 °C. Under these conditions, FUS RRM exchanges between folded and unfolded states in slow exchange on the NMR chemical shift timescale to produce two sets of NMR resonances which are sufficiently resolved in 3D HNCO experiments, in which peaks are separated according to the ^1^H and ^15^N chemical shifts of a given residue and the ^13^CO shift of the preceding amino acid(27). Interactions between proximal FUS RRM molecules are predicted to increase as a function of protein concentration. This leads to decreased peak intensities for residues at the sites of contact relative to peaks derived from regions with no interactions because the decreased dynamics at contact points and/or conformational heterogeneity at these sites results in a faster decay of NMR signals. Interactions can, therefore, be quantified by monitoring the decrease in HNCO peak intensities in the high (*I*_*concentrated*_) *vs*. low (*I*_*dilute*_) concentration datasets, after adjusting for the 4-fold concentration disparity between samples. Thus, *I*_*concentrated*_ */ I*_*dilute*_ ratios report on sites of transient intermolecular association, *i*.*e*., interaction hotspots (Fig. 2A). As separate sets of peaks are observed for folded and unfolded conformers of FUS RRM, the magnitude of the intermolecular interactions (intensity losses) for each state can be probed separately, as shown for S282, G291 and T338 (Fig. 2B). Little intensity change is observed for any of the FUS RRM peaks derived from the folded conformer (Fig. 2B, *top row*), with significant intensity losses evident for peaks from G291 and T338 of the unfolded state, but not for S282, for example, in a comparison of high *vs*. low concentration datasets (Fig. 2B, *bottom row*). Therefore, residues G291 and T338 in the unfolded state of FUS RRM form transient intermolecular contacts or are part of a region that does.

**Figure 2.**
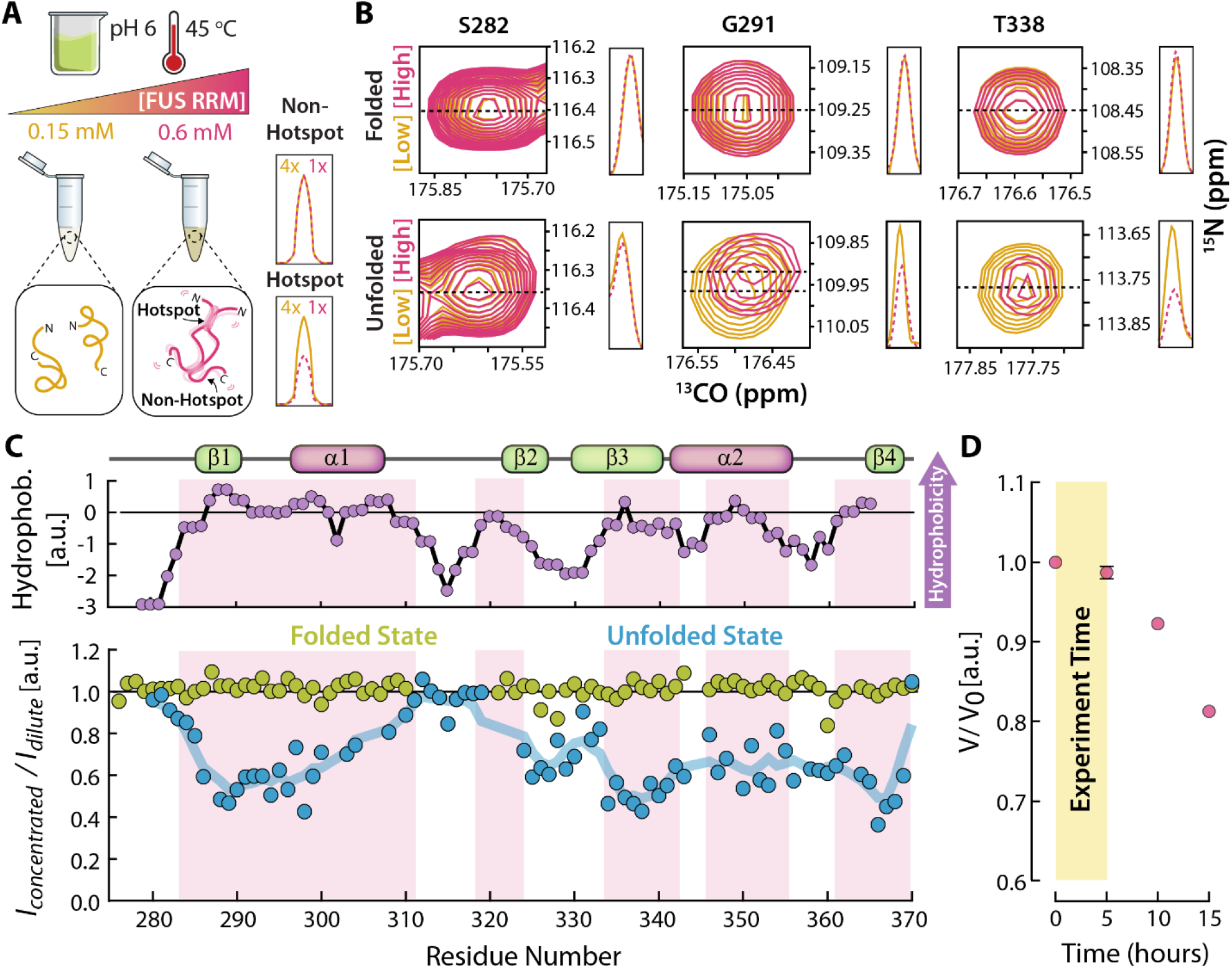
Intermolecular FUS RRM interactions. (**A**) Cartoon illustrating the use of concentration-dependent (0.15 mM, solid yellow, *vs*. 0.6 mM, dashed red) peak intensity measurements to identify intermolecular interactions driving FUS RRM aggregation. A comparison of NMR peaks derived from ‘hotspot’ and ‘non-hot-spot’ regions under conditions of high and low protein concentrations is shown. Peak intensities in 3D HNCO datasets are scaled (4x) to adjust for the 4-fold concentration disparity between samples, so that peak intensity differences reflect intermolecular association. (**B**) Representative peaks derived from folded and unfolded conformations of FUS RRM in low (0.15 mM, *yellow*) and high (0.6 mM, *red*) concentration samples, 45 °C. Slices through the ^13^CO dimension are shown for straightforward interpretation of intensity changes. (**C**) Hydrophobicity of the FUS RRM sequence (*top, purple symbols*), as determined using the Kyte and Doolittle hydrophobicity scale(41). HNCO peak intensity ratios of FUS RRM peaks in concentrated (0.6 mM) *vs*. dilute (0.15 mM) samples (*bottom*), normalized for concentration differences. Pink highlights indicate regions of high hydrophobicity. The blue line shows a rolling average of the unfolded state intensity profile, where the intensity ratios for a given residue and its four adjacent neighbors (two on either side) are averaged. (**D**) Time-dependent losses of all NMR-visible FUS RRM amide signals in 1D ^15^N,^13^C-edited (HNCO-based) ^1^H NMR spectra indicate that, under these conditions, intermolecular interactions occur without significant aggregation on the timescale of the measurements (*yellow highlighted region*). Nonetheless, these interactions promote aggregation over longer time periods (> 5 hours).

Overall, intermolecular interactions between folded FUS RRM conformers are not observed, with *I*_*concentrated*_ */ I*_*dilute*_ values close to one for the majority of the sequence (Fig. 2C, *bottom, green symbols*). In contrast, intermolecular contacts are observed for significant regions of FUS RRM molecules in the unfolded state, including much of the C-terminus extending from approximately G335-A369 and another segment spanning I287-I308 (Fig. 2C, *bottom, blue symbols*). Yet a third segment, L324-T330, shows somewhat weaker contacts. The interaction hotspots are enriched in hydrophobic amino acids (Fig. 2C, *red highlighted regions*), whereas negligible intermolecular contacts are observed for the ^312^KTNKKTG^318^ hydrophilic stretch. It is noteworthy that FUS RRM is stable for the duration of the experiment (i.e., does not form NMR invisible aggregates; Fig. 2D, *yellow highlighted region*). Nonetheless, global intensity losses are observed over longer timeframes (Fig. 2D, *non-highlighted region*) indicating that the interactions between unfolded FUS RRM chains mapped here are involved in aggregation.

### CAPRIN1 condensates promote FUS RRM unfolding yet prevent aggregation

Our NMR experiments established that aggregation of FUS RRM is suppressed in mixed solutions of 0.5 mM FUS RRM and 4 mM CAPRIN1 (Fig. 1C). We were interested in extending these studies to a FUS RRM:CAPRIN1 condensate, initially to understand how the FUS RRM conformational landscape is affected in the condensed phase, as well as to evaluate whether aggregation would be prevented in this proteinaceous, highly concentrated environment. To this end we prepared ^2^H,^15^N,^13^C-ILV labeled FUS RRM (only methyl groups of Ile δ1, Leu δ1,δ2, and Val γ1,γ2 are NMR observable, ^13^CH3, with only one of the two prochiral methyl groups of Leu and Val ^13^CH3-labeled(18)) and mixed it with ^1^H,^14^N,^12^C CAPRIN1 in pH 6 buffer. As we have done previously, the Leu and Val residues in CAPRIN1 (3 in total; there are no Ile) were replaced so as not to interfere with LV-methyl signals from the client(21). Phase separation of the mixture was induced through the addition of NaCl to a final concentration of 200 mM, as described in *SI Appendix*. The resulting phase-separated solution was transferred into a 3 mm NMR tube (Fig. 3A) with the condensed phase covering the entirety of the receiver coil, ensuring that the NMR signals are solely derived from it. After completion of NMR experiments on the condensed phase, a portion of the dilute phase was decanted from the top and placed into a second NMR tube so that it too could be studied.

**Figure 3.**
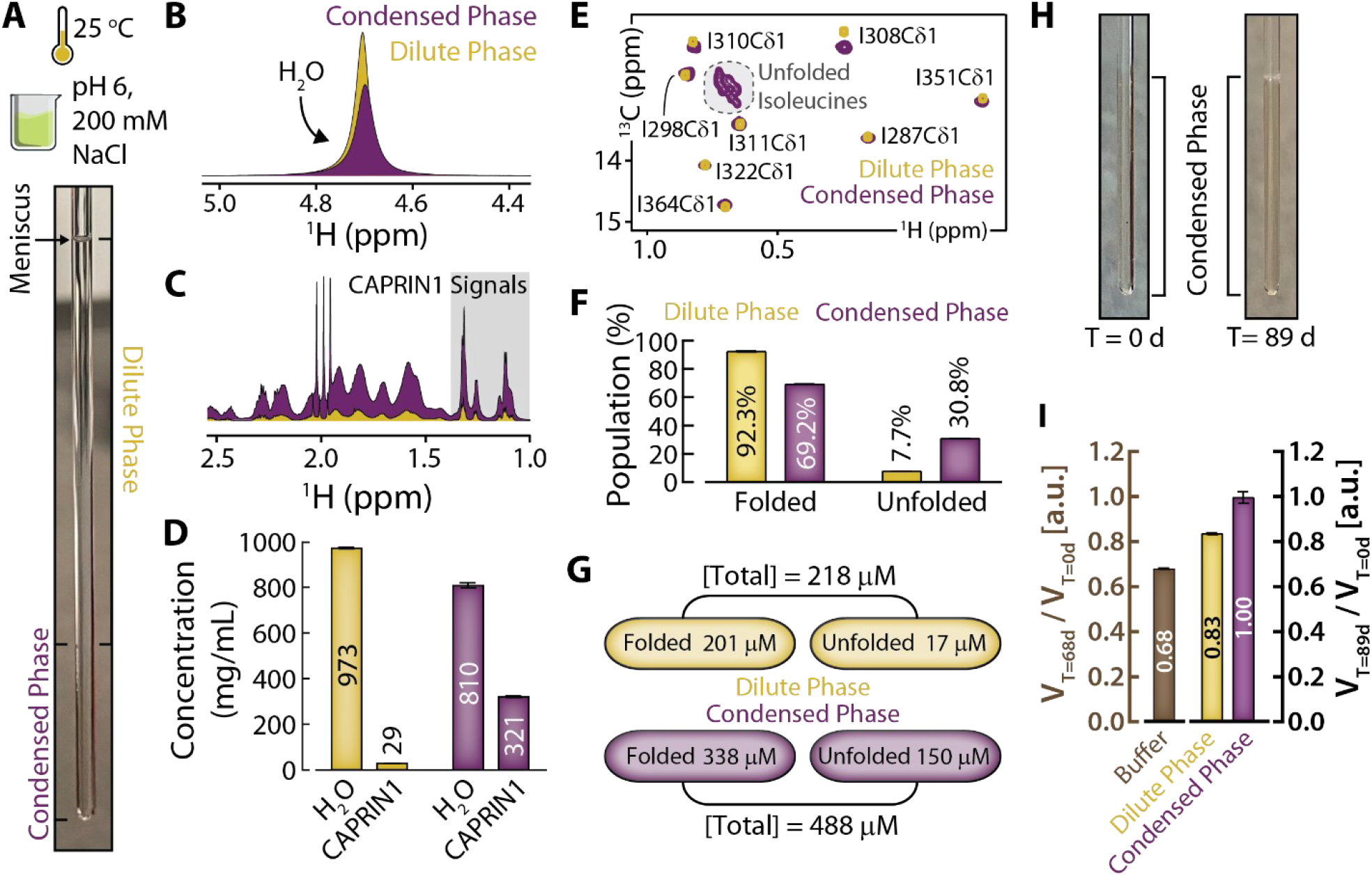
CAPRIN1 condensates promote unfolding and suppress aggregation of FUS RRM. (**A**) Phase-separated ^2^H,^15^N,^13^C-ILV FUS RRM:^1^H,^14^N,^12^C CAPRIN1 NMR sample with a black marker indicating the dilute and condensed phase boundary and an arrow showing the solution-air boundary (meniscus). The phase-separated sample was prepared in 20 mM MES pH 6 buffer containing 200 mM NaCl. All spectra were recorded at 25 °C on an 800 MHz spectrometer. Only the condensed phase is observed, as the NMR detection coil covers only this region. (**B-C**) ^1^H NMR spectra used to quantify the amount of (**B**) H^2^O and (**C**) CAPRIN1 in the dilute (*yellow*) and condensed (*purple*) phases of FUS RRM:CAPRIN1. (**D**) Quantification of the [H^2^O] and [CAPRIN1] in dilute (*yellow*) and condensed (*purple*) phases, based on the integrals of the peaks shown in (**B-C**) relative to peaks in spectra recorded on reference samples with known concentrations. Note the dilute and condensed phase spectra were recorded using separate samples. (**E**) ^1^H,^13^C delayed decoupling methyl-TROSY spectra of ^2^H,^15^N,^13^C-ILV FUS RRM in dilute (*yellow*) and condensed (*purple*) phases. Signals derived from unfolded FUS conformers are indicated by the grey highlighted regions. (**F**) Populations of folded and unfolded FUS RRM in the dilute (*yellow*) and condensed (*purple*) phases of FUS RRM:CAPRIN1, 25 °C, based on the volumes of Ile peaks derived from folded and unfolded conformers shown in (**E**), taking into account differences in transverse relaxation between NMR signals in each state. The total volume of the Ile region from the unfolded state was determined using a sum-over-box approach. Errors were determined from the propagation of uncertainties derived from the peak fitting analysis. (**G**) Concentrations of total FUS, as well as folded and unfolded FUS RRM states in the dilute (*yellow*) and condensed (*purple*) phases of CAPRIN1, as determined through comparison of peak volumes with a reference state of known concentration. (**H**) Phase-separated ^1^H,^15^N,^13^C FUS RRM: ^1^H,^14^N,^12^C CAPRIN1 samples at T = 0 days (*left*) and T = 89 days (*right*), highlighting the lack of visible aggregates. (**I**) Relative integrals of FUS RRM signals at T = 89 d *vs*. T = 0 d in the dilute (*yellow*) and condensed (*purple*) phases. The ratio of integrals of FUS RRM signals at T = 68 d *vs*. T = 0 d in buffer with a concentration equivalent to that in the condensed phase is also shown (*brown*). The error bars are determined based on the signal-to-noise ratio in the spectra.

We measured concentrations of water and CAPRIN1 in the condensed phase (25 °C), as well as in the dilute phase for comparison, to understand the contributions of each to the solvation of the client FUS RRM. By comparing peak volumes of the H_2_O signal in ^1^H NMR spectra recorded of each of the phases with a spectrum from a reference that matched the buffer and salt composition of the phase-separated sample, H_2_O concentrations of ∼ 973 mg/mL and ∼ 810 mg/mL were calculated for dilute and condensed environments, respectively (Fig. 3B). Similarly, concentrations of CAPRIN1 were assessed by focusing on a region of the ^1^H spectrum containing CAPRIN1 signals exclusively (1-1.4 ppm, Fig. 3C) *i*.*e*., excluding those signals arising from ^2^H,ILV-labeled FUS RRM (upfield of 1 ppm). These measurements establish that CAPRIN1 significantly contributes to the solvation of FUS RRM in the condensed phase (H_2_O ∼ 810 mg/mL (45 M) *vs*. CAPRIN1 321 mg/mL (30 mM); Fig. 3D).

To assess how the dilute and condensed phase solvent environments impact the conformational landscape of FUS RRM, we recorded ^1^H,^13^C delayed decoupled HMQC spectra(20) (ddHMQC). Coherence transfer selection gradients were used to effectively suppress the intense CAPRIN1 scaffold signals, which would otherwise obscure signals derived from the FUS RRM client, as described previously(21). These methyl-TROSY(19) based experiments are crucial for mapping protein conformational landscapes in highly viscous solutions, such as those presented by the proteinaceous condensed phase environment(14, 21). In previous studies focused on either SOD1 or RNA clients, methyl-based spectra were essential, as amide correlations were not observed in spectra of condensates under the conditions used for the measurements(7, 21). ddHMQC spectra recorded on the dilute and condensed phase samples of FUS RRM:CAPRIN1 revealed two separate sets of FUS RRM resonances in slow-exchange on the NMR chemical shift timescale that are derived from folded and unfolded FUS RRM conformations (Fig. 3E). This includes eight well-dispersed isoleucine resonances from folded FUS RRM and another set of overlapping signals arising from the unfolded ensemble (Fig. 3E, *grey shaded region*). Populations of the two states were quantified from the isoleucine peak volumes in fully relaxed ddHMQC spectra shown in Fig. 3E. Peak volumes were corrected for losses resulting from transverse relaxation during the fixed delays of the experiment, allowing for accurate population measurements(28). Notably, we observe a significant reduction in the population of the folded FUS RRM, from 92% in the dilute phase to 69% in the condensed phase, 25 °C (Fig. 3F), corresponding to a difference in unfolding free energies, 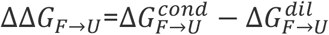 of -4.1 kJ/mol (*SI Appendix*, Table S1). By comparing peak volumes in methyl-HMQC spectra in each of the two phases to those in a reference spectrum with a known concentration of FUS RRM, concentrations of folded and unfolded conformers in dilute and condensed phases were obtained (Fig. 3G; *SI Appendix*, Materials and Methods). A partition coefficient, *K*_*P*_, (total [FUS RRM] in condensed phase divided by total [FUS RRM] in dilute phase) of 2.2 was calculated, indicating that FUS RRM has a mild preference for the condensed phase solvent environment. Under these non-stress conditions, the folded FUS RRM remains the major conformation in the condensed phase (69%). However, under mild heat stress (40 °C) the folded conformer population decreases to 36% in the condensed phase (*SI Appendix*, Table S1).

Given that the unfolding of FUS RRM under acidic and moderate heat stress conditions drives its aggregation (Fig. 2C,D) and that the CAPRIN1 solvent environment significantly increases the population of the client unfolded ensemble (Fig. 3F,G), we wondered whether FUS RRM aggregation would be promoted in CAPRIN1 condensates. To investigate this, we monitored amide intensities of FUS RRM in spectra recorded of dilute and condensed phase samples over approximately 3 months (Fig. 3H). A sample of similarly labeled FUS RRM dissolved in buffer, and with a total FUS RRM concentration matching that in the condensed phase (∼0.5 mM), was also monitored for comparison. No significant change in the FUS RRM signal was observed for the condensed phase (∼30 mM CAPRIN1), while decreases of approximately 20% and 40% were noted for the dilute phase (with a substantially lower CAPRIN1 concentration of ∼2.6 mM) and the buffer sample (lacking CAPRIN1), respectively (Fig. 3I). Thus, the rich CAPRIN1 environment of the condensate solvates the unfolded ensemble of FUS RRM and prevents formation of higher molecular weight aggregates.

### Unambiguous detection of unfolded FUS RRM amide signals in the condensed phase

To understand how the CAPRIN1 scaffold stabilizes the unfolded form of FUS RRM it is necessary to map scaffold interactions to specific sites on the client. This process is challenged by the fact that the abundancies of ^14^N and ^15^N isotopes in unenriched nitrogen are 99.7% and 0.3%, respectively. Thus, at a concentration of 30 mM CAPRIN1 in the condensed phase, the effective concentration of ^15^N at a given backbone position in CAPRIN1 is approximately 90 µM, which is not negligible in comparison to the concentration of the unfolded FUS RRM (∼150 µM at 25°C, ∼335 µM at 40°C) (*SI Appendix*, Table S1). The resulting overlap of intrinsically disordered CAPRIN1 and unfolded FUS RRM signals precludes obtaining unambiguous site-specific information on client interaction sites (Fig. 4A), even when measurements are carried out at 40°C where there is almost a four-fold difference in relative NMR active concentrations of scaffold and client proteins (*SI Appendix*, Table S1). Therefore, we prepared CAPRIN1 in minimal media supplemented with 99.99% ^14^N-(NH_4_)_2_SO_4_ as the sole nitrogen source, decreasing the NMR active scaffold concentration to 3 µM. The large discrepancy in concentration between ^15^N-unfolded FUS RRM *vs*. ^15^N-CAPRIN1 that results using this labeling scheme now ensures the selective observation of unfolded FUS RRM amide signals, as shown in Fig. 4A (*red vs. blue*). The zoomed-in expansions shown in Figs. 4B-D make it clear that without ^15^N-depleted scaffold the majority of unfolded FUS RRM amide signals could not be distinguished from those of CAPRIN1 and quantitative analysis of scaffold interactions sites would not be possible.

**Figure 4.**
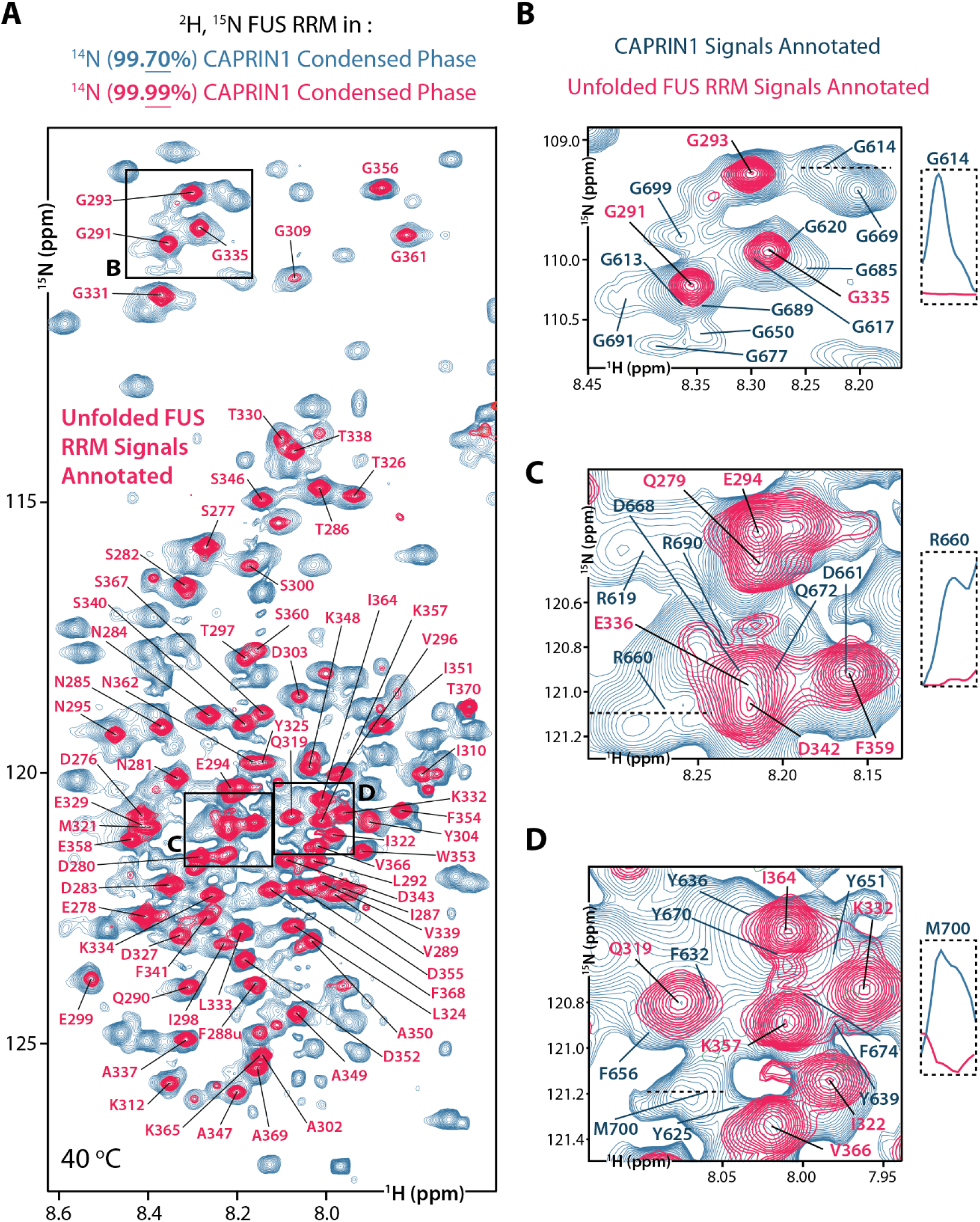
Selective labeling enables background-free observation of condensed phase FUS RRM signals from unfolded conformers in ^1^H,^15^N spectra. (**A**) Overlay of ^1^H,^15^N TROSY-HSQC spectra recorded on ^2^H,^15^N FUS RRM:CAPRIN1 condensed phase samples prepared with either 99.7% ^14^N CAPRIN1 (*blue*) or 99.99% ^14^N CAPRIN1 (*red*; *Materials and Methods*). Spectra were recorded at 40 °C, 800 MHz. Unfolded FUS RRM signals are annotated in red. (**B-D**) Zoomed-in expansions of spectral regions shown in (**A**). Signals from the unfolded FUS RRM and from CAPRIN1 are annotated in red and blue, respectively. Slices through the ^1^H dimension are shown at specified positions (*dashed lines*) to indicate the absence of CAPRIN1 signals in samples prepared with 99.99% ^14^N CAPRIN1.

### Mapping heterotypic unfolded client:scaffold interaction sites on the client

Having established a labeling strategy for selectively recording amide signals from the unfolded FUS RRM (Fig. 4), we next focused on quantifying intermolecular interactions with CAPRIN1. To this end we recorded a Nuclear Overhauser Effect (NOE) based-experiment in which magnetization is transferred from protons coupled to ^13^C (^13C^H) in CAPRIN1 (i.e., aliphatic and aromatic protons) to protons attached to ^15^N (^15N^H) in FUS RRM (Fig. 5A, *SI Appendix*, Fig. S2). To ensure that NOEs arise exclusively from intermolecular interactions between CAPRIN1 and FUS RRM chains we prepared a condensed phase sample consisting of ^2^H,^12^C,^15^N FUS RRM and ^1^H,^14^N (99.99%) CAPRIN1, where one-third of the CAPRIN1 molecules were also ^13^C-labeled. Datasets were recorded as 2D ^1^H-^15^N TROSY-HSQC spectra with NOEs ‘read-out’ on the amides of FUS RRM (*SI Appendix*), as described in detail in our previous work focusing on measurement of homotypic scaffold interactions within a condensed CAPRIN1 sample(22). Importantly, the low natural abundance of ^13^C (1.1%) coupled with deuteration ensures that intramolecular NOEs within a ^2^H,^12^C,^15^N FUS RRM chain are not observed. This is demonstrated for a 0.5 mM ^2^H,^12^C,^15^N FUS RRM sample in buffer (*SI Appendix*, Fig. S3).

**Figure 5.**
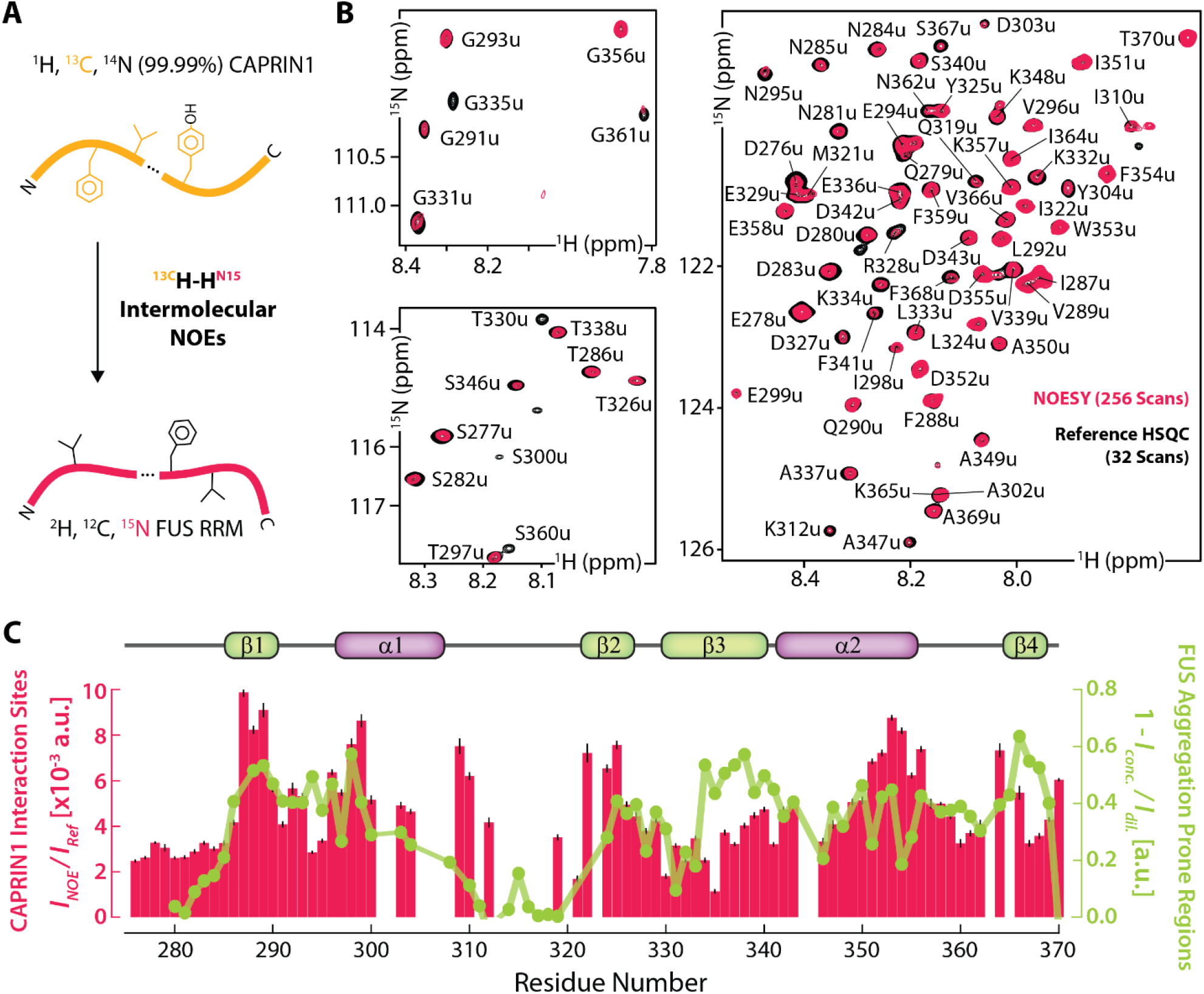
CAPRIN1 interaction sites in FUS RRM overlap with FUS RRM aggregation-prone regions. (**A**) Schematic of the intermolecular NOE experiment quantifying magnetization transfer from protons attached to ^13^C (*orange chain*) to protons bonded to ^15^N (*red chain*) on separate chains. Note that the labeling scheme ensures that NOEs can only arise from interactions between CAPRIN1 and FUS RRM (*SI Appendix*, Figure S3). (**B**) Expanded regions from experiment schematically illustrated in A, recorded with a mixing time of 250 ms. The NOESY spectrum (*red*) is superimposed onto the corresponding regions from a ^1^H,^15^N TROSY-HSQC spectrum (*black*); both data sets are recorded at 1 GHz, 40 °C. Note that the NOE experiment is recorded with 8-fold more scans than the HSQC. Only NOEs to the unfolded state of FUS RRM were observed; these are annotated. (**C**) Intermolecular NOE profile (*red bars*) after normalizing NOE peak intensities to the intensities of the corresponding amide correlations in an HSQC-TROSY dataset. In addition, the data were normalized to account for the differences in the numbers of scans in the two experiments (i.e., intensities of the NOE data set were divided by 8 = 256/32). Errors were estimated based on the propagation of uncertainties derived from the peak fitting analysis. The unfolded FUS RRM aggregation profile shown in Fig. 2C is overlaid in green (*circles and connected line*), rearranged so that 1 – (*I*_*conc*.._ / *I*_*dil*._) is shown. Residues lacking data are those for which resonance assignments are not available.

To maximize the signal-to-noise ratios of the NOE correlations we recorded spectra at 40°C, where the concentration of unfolded FUS RRM is two-fold higher than at 25°C (SI A*ppendix*, Table S1). Figure 5B shows expanded regions from the NOESY dataset (*red*, 250 ms mixing time), overlaid onto corresponding regions from a reference TROSY-HSQC spectrum (*black*). In general, peak intensities in the NOESY spectrum increase as the interactions between CAPRIN1 and FUS RRM become stronger, corresponding to closer client/scaffold distances and/or increases in lifetimes of interaction. A plot of the intermolecular NOE intensities as a function of FUS RRM residue number is shown in Figure 5C (*red*). Residue-specific differences in peak intensities arising from signal loses due to relaxation during delays in the experiment and/or solvent exchange were taken into account through normalization of NOEs with peak intensities from a TROSY-HSQC dataset recorded with identical parameters as for the NOE experiment. Intermolecular NOEs (*red*) are observed throughout the unfolded FUS RRM sequence (Fig. 5C), consistent with the overall solvation of unfolded FUS RRM conformers by CAPRIN1. Nonetheless, several regions with stronger intermolecular contacts are observed, including ^287^IFVQ^290, 296^VTIES^300, 322^INLY^325^ and ^351^IDWFDG^356^. These regions include aromatic and negatively charged residues, that would naturally interact with aromatic- and arginine-rich segments of CAPRIN1. Notably, sites of heterotypic FUS RRM:CAPRIN1 interactions (Fig. 5C, *red*) significantly overlap with regions of homotypic FUS RRM-FUS RRM contacts (Fig. 5C, *green*), suggesting that CAPRIN1 solvation of unfolded FUS RRM inhibits interactions between protomers that promote aggregation.

## DISCUSSION

Phase separation generates two or more co-existing phases distinguished by differences in molecular composition. The asymmetric partitioning of various molecules, such as ions, metabolites, nucleic acids, and proteins across these phases gives rise to emergent properties that shape condensate function. These include distinct pH(29, 30) and dielectric values(31), as well as material(32) and solvent characteristics(33, 34) that collectively modulate the conformational landscapes of nucleic acids and proteins and, in turn, their function. Indeed, we and others have shown that condensates can alter the thermodynamics and kinetics of nucleic acid hybridization(7, 35), shift the folding-unfolding equilibria of client proteins(14, 36, 37), enable access to protein conformational states that evolve into aberrant aggregates(14), and effect the rates of chemical reactions(5, 6). Sequestration of proteins into condensates can also locally increase their concentration, promoting nucleation and aggregation events(9, 11). Nonetheless, several studies have demonstrated that condensates can also suppress client aggregation, functioning as protective reservoirs for aggregation-prone client proteins(16, 17). Important goals in this context are to elucidate the specific interactions between client proteins and condensate scaffolding molecules that inhibit aggregation in these cases, and to establish how the conformational landscapes of client proteins are modulated to achieve this effect.

Here, we have applied solution NMR spectroscopy to investigate the aggregation behavior of the RRM from the stress granule client protein FUS, both outside and within model condensates composed of a C-terminal disordered region from the stress granule scaffold protein, CAPRIN1. We find that FUS RRM aggregates under acidic conditions and elevated temperatures commonly associated with cellular stress (Fig. 1C), and that aggregation is mediated through access to an unfolded state of the protein (Fig. 2 and Fig. 6, *top*). Our concentration-dependent NMR studies identify regions of transient intermolecular contacts between unfolded FUS RRM chains, which include the hydrophobic segments spanning I287-I308 and G335-A369, whereas negligible interactions are found for the ^312^KTNKKTG^318^ hydrophilic stretch and between folded FUS RRM conformers (Fig. 2). These interactions facilitate aggregation of FUS RRM in buffer and, to a lesser extent, in the dilute phase of phase-separated FUS RRM:CAPRIN1 solutions, where the concentration of CAPRIN1 is approximately 3 mM (Fig. 1C, 3I). However, no aggregation was observed in the condensed phase (30 mM CAPRIN1), despite an approximate 2-fold increase in the total FUS RRM concentration relative to the dilute phase (Fig. 3G,3H and *SI Appendix*, Table S1), a significant increase in the fractional population of the unfolded RRM client, and close to an order of magnitude increase in the absolute concentration of the unfolded ensemble at 25°C (Fig. 3E-G and *SI Appendix*, Table S1). Insights into the protective role of CAPRIN1 can be obtained at atomic resolution by recording intermolecular NOEs in the condensed phase. Our results indicate that there are CAPRIN1 interaction sites throughout the unfolded FUS RRM, including several that are of particular significance, such as ^287^IFVQ^290, 296^VTIES^300, 322^INLY^325^ and ^351^IDWFDG^356^ (Fig. 5C, *red* and Fig. 6, *bottom*). Notably, these regions largely overlap with sites of homotypic FUS RRM-FUS RRM interactions (Fig. 5C, green), indicating that the high concentration of the CAPRIN1 “solvent” may outcompete the interactions between unfolded FUS RRM protomers that otherwise would drive aggregation.

**Figure 6.**
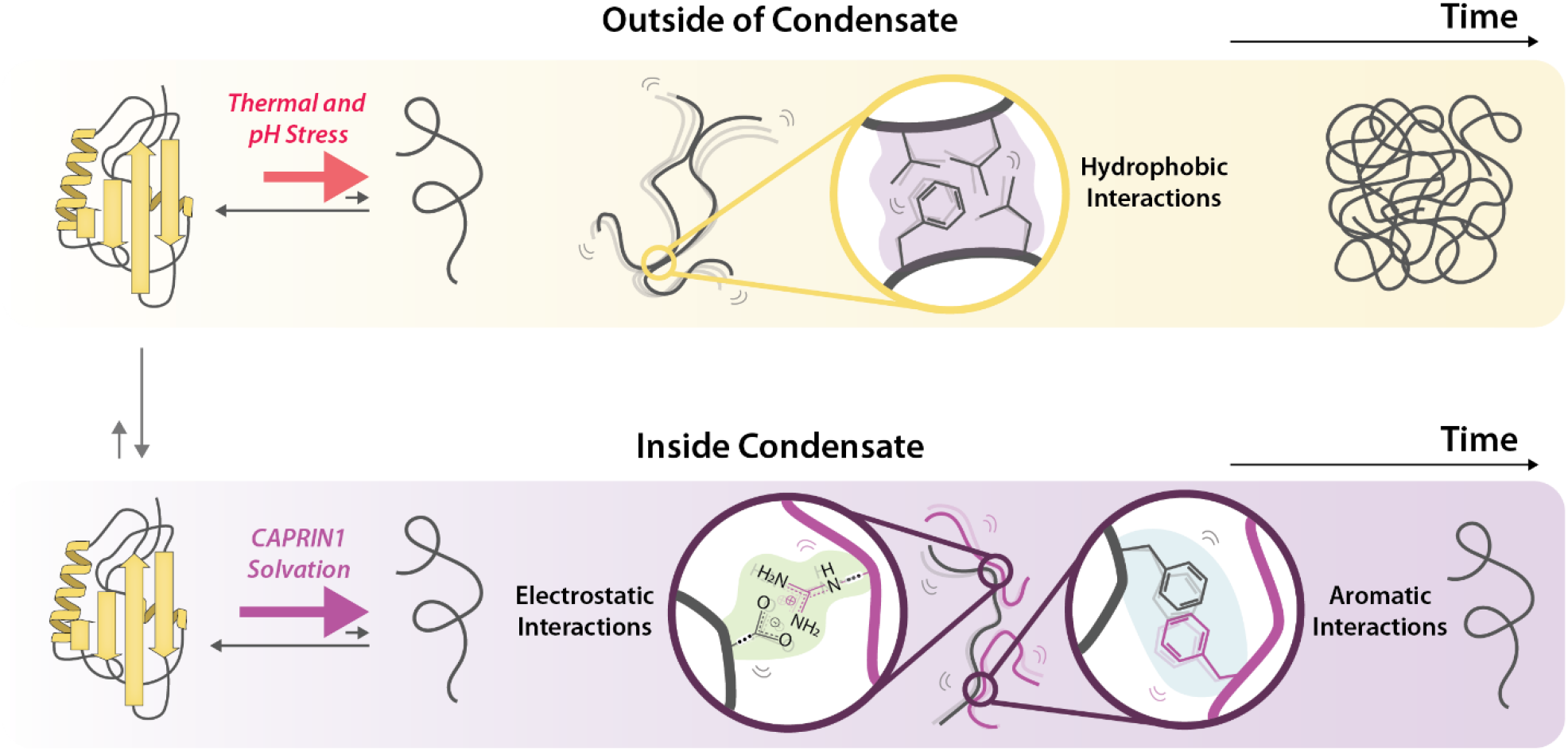
Schematic model highlighting how CAPRIN1 condensates suppress FUS RRM aggregation. Homotypic interactions between unfolded FUS RRM client molecules driving aggregation (*top*) and the heterotypic FUS RRM:CAPRIN1 interactions suppressing aggregation in CAPRIN1 scaffold condensates (*bottom*). Our NMR studies indicate that thermal and pH stresses unfold FUS RRM (*top, left*), enabling hydrophobic interactions (*top, zoomed-in expansion*) between unfolded FUS RRM chains that promote aggregation (*top, right*). Partitioning of the FUS RRM client into CAPRIN1 scaffold condensates promotes FUS RRM unfolding (*bottom, left*) without causing aggregation (*bottom, right*), as heterotypic electrostatic and aromatic interactions between FUS RRM and CAPRIN1 (bottom, *zoomed-in expansions*) outcompete homotypic intermolecular interactions between unfolded FUS RRM chains.

A large body of work suggests that condensates can promote nucleation and aggregation of proteins, including scaffolds that assemble the condensate(9, 12) and clients that partition into condensates formed by other components(17, 38). Our results, however, establish that for the FUS RRM:CAPRIN1 condensed phase studied here, the condensate scaffold molecules shield client interprotomer homotypic interactions, thereby delaying or completely suppressing aggregation. These findings are consistent with and complementary to results from a study exploring the aggregation kinetics of the Alzheimer’s disease-associated peptide, amyloid-beta 42 (Aβ42), in condensates formed by a designer scaffold comprised of adenylate kinase (AK) conjugated at the N- and C-termini with the low complexity domain of LAF-1 *i*.*e*. Laf1-AK-Laf1(16). The authors showed that Aβ42 aggregation is inhibited in the dilute phase due to sequestration of Aβ42 inside the condensate, while interactions between the scaffold molecules and the Aβ42 client suppress aggregation in the condensed phase. Notably, they find that addition of sub-stoichiometric (micromolar) amounts of the scaffold is sufficient to attenuate aggregation, indicating specific and protective bimolecular client-scaffold interactions. In contrast, in our study we observe that high millimolar concentrations of the CAPRIN1 scaffold, such as those found inside the FUS RRM:CAPRIN1 condensed phase (30 mM), are required for suppressing aggregation (Fig. 3I, *condensed vs. dilute phase*). This reinforces the notion that low-affinity interactions of a proteinaceous solvent can have profound impacts on the conformational landscape of clients(39).

Another study demonstrated that droplets formed by binary mixtures of polyanions and polycations suppress aggregation of the Parkinson’s disease-associated alpha synuclein protein(17). The authors showed that these model condensates can inhibit multiple nucleation steps of the aggregation cascade, including primary nucleation that controls the lag time for aggregation as well as elongation and secondary nucleation events that regulate the growth rate of fibrils. Based on these observations, the authors speculate that inhibition of aggregation arises from condensate scaffolds interacting with and stabilizing the monomeric form of the client. Our work provides direct evidence supporting this model by showing, at atomic resolution, a strong correspondence between the homotypic client-client interactions driving aggregation and the heterotypic client-scaffold interactions that prevent it.

In this regard, our work underscores the unique contribution that solution NMR spectroscopy can provide for obtaining atomic resolution insights into client-scaffold interactions and in mapping the free energy landscapes of client molecules within condensates. Notably, while other widely applied techniques, such as single molecule fluorescence resonance energy transfer, have offered critical insights into the global conformations of biomolecules in condensates or the strength of their interactions(40), they cannot easily distinguish how different conformers of a client protein, *e*.*g*., folded and unfolded states, interact with neighboring molecules. As we have highlighted in this work, solution NMR can not only distinguish between various protein conformations but also map, at atomic resolution, their interactions. Therefore, NMR offers a more nuanced understanding of the complexity of molecular interactions in phase-separated systems. The labeling approaches and NMR methods presented here are likely to be applicable to other client-scaffold systems. The insights gained from these studies will provide a more comprehensive understanding of the mechanisms by which the conformational landscapes of biomolecules are modulated within the condensed, proteinaceous environment of a phase-separated system, and, hence, how their functions can be regulated by it.

## Acknowledgements

R.A. acknowledges post-doctoral support from the Hospital for Sick Children and the Canadian Institutes of Health Research (CIHR). J.D.F.-K. acknowledges support from the CIHR (PJT-190060), the Natural Sciences and Engineering Council of Canada (NSERC) (2024-05725), and from the Canada Research Chairs Program. L.E.K. acknowledges support from the CIHR (FND-503573) and NSERC (2024-03872). CD data were acquired using instrumentation at the Structural Biophysical Core Facility (Hospital for Sick Children). We thank Philip Rößler for helpful discussions on the use of ^15^N-depleted ammonium sulfate for selective unlabeling of proteins and Professor Alvar D. Gossert, ETH, Zurich, for kindly providing some initial 99.99% ^14^N-(NH_4_)_2_SO_4_ material. All data within the manuscript, including NMR pulse programs and parametersets, will be provided on Zenodo.

