## Supporting Information for "Client-scaffold interactions suppress aggregation of a client protein in model condensates"

### Materials and Methods

#### Expression and purification of CAPRIN1, and RRMs from FUS, TAF-15, and EWSR1

The C-terminal low-complexity disordered region of human CAPRIN1 comprising residues Ser607-Gln707 and containing the mutations N623T, N630T, V610A, L621A, referred to henceforth as CAPRIN1, was subcloned into a pET-His-SUMO vector. Substitution of Thr for Asn at positions 623 and 630 eliminates the formation of Iso-Asp linkages that would otherwise occur over time, as discussed previously(42), while removal of Val and Leu residues in the scaffold enables observation of methyl signals from spectra of the FUS RRM client without interference from peaks derived from CAPRIN1(21). *Escherichia coli* BL21 (DE3) RIPL cells were transformed with the vector bearing the CAPRIN1 sequence and grown to an $OD_{600}$ ~ 0.6 – 0.8. Protein expression was induced with 0.5 mM IPTG and allowed to continue overnight at 25 °C in LB (M9 minimal media) for unlabeled (isotopically labeled) protein. For production of 99.99% $^{14}N$ CAPRIN1, cells were supplemented with 1 g/L 99.99% $^{14}N$-$(NH_4)_2SO_4$ (Sigma Aldrich) as the sole nitrogen source. Uniformly $^{13}C$-labeled CAPRIN1 was produced by supplementing M9 minimal media with 3 g/L D-Glucose-13$C_6$ (Cambridge Isotope Laboratories) as the sole carbon source. Cells were harvested via centrifugation, resuspended in lysis buffer (6 M guanidinium chloride, 50 mM Tris pH 8, 500 mM NaCl, 10 mM Imidazole), and sonicated for 15 minutes (2 s on, 2 s off). Lysed cells were spun down at 13,800 g for 1 hour, and the supernatant was loaded onto a Ni-NTA column (GE Healthcare) equilibrated with lysis buffer. The column was washed extensively with lysis buffer and CAPRIN1 was eluted with a solution of 50 mM Tris pH 8, 150 mM NaCl, 400 mM imidazole. The His-SUMO tag was cleaved with HisSUMO protease while exchanging against 50 mM Tris pH 8, 150 mM NaCl, 10 mM imidazole, 2 mM β-mercaptoethanol buffer at 4 °C. The cleaved protein was loaded onto a Ni-NTA column to remove the His-SUMO tag and HisSUMO protease, concentrated, and injected onto a Superdex75 (26/600) column equilibrated with 3 M guanidinium chloride, 50 mM Tris pH 8. The purified protein fractions were pooled and stored at -20 °C until use.

The human FUS RRM (D276-T370), TAF-15 RRM (D232-E323), and EWSR1 RRM (D353-S453) sequences all bearing an N-terminal diglycine (GG) were subcloned into pET-His-SUMO vectors and transformed into *Escherichia coli* BL21 (DE3) RIPL cells. The cells were grown to an $OD_{600}$ ~ 0.6-0.8, protein expression induced with 0.5 mM IPTG, and protein expression continued overnight at 18 °C in M9 $D_2O$ ($H_2O$) media supplemented with 3 g/L $d_7$-glucose (glucose) for deuterated (non-deuterated) protein production. Uniform $^{15}N$ isotopic labeling was achieved by the addition of 1 g/L $^{15}N$ ammonium chloride to the M9 growth medium. Methyl labeling was achieved by the addition of 100 mg/L of 2-keto-3-methyl-$d_3$-3-$d_1$-4-$^{13}C$-butyrate (for non-stereospecific labeling of Leu, Val-$^{13}CH_3$/$^{12}CD_3$) and 60 mg/L of 2-keto-3-$d_2$-4-$^{13}C$-butyrate (for $^{13}CH_3$ labeling of Ileδ1) one hour prior to the induction of protein expression. Cells were harvested by centrifugation, resuspended in lysis buffer (50 mM Tris pH 8, 300 mM NaCl, and 20 mM Imidazole) supplemented with lysozyme, RNAse A (Roche) and protease inhibitor cocktail tablets (Roche), and sonicated for 15 minutes (2 s on, 2 s off). Lysed cells were spun down at 13,800 g for 1 hour, and the supernatant was loaded onto a Ni-NTA column (GE Healthcare). After extensive washing with lysis buffer, the RRMs were eluted with a buffer comprised of 50 mM Tris pH 8, 150 mM NaCl, and 400 mM Imidazole. The His-SUMO tag was cleaved with HisSUMO protease during constant buffer exchange with 50 mM Tris pH 8, 150 mM NaCl, 20 mM imidazole, 2 mM β-mercaptoethanol, 10% glycerol buffer at 4 °C. The cleaved protein was isolated through another round of Ni-NTA purification, concentrated, and loaded onto a Superdex75 (26/600) column equilibrated with 50 mM Tris pH 8. The pure protein-containing fractions were pooled and stored at 4 °C until use.

**Preparation of FUS RRM:CAPRIN1 condensed phases**

CAPRIN1 and FUS RRM were buffer-exchanged into a solution of 20 mM MES pH 6.0, 0.5 mM EDTA using a HiPrep 26,10 Desalting column. Concentrated stocks of FUS RRM (> 1 mM) and CAPRIN1 (> 8 mM) were mixed on ice and allowed to equilibrate for 30 minutes. Phase separation was induced through 10% dilution of the FUS RRM:CAPRIN1 mixture with 20 mM MES pH 6.0, 200 mM NaCl, 0.5 mM EDTA, 100% $D_2O$, resulting in a final NaCl concentration of 200 mM and 10% $D_2O$. A portion of the phase-separated mixture was then transferred into a 3-mm NMR tube and stored at 4 °C until the phase-separated droplets coalesced into a homogeneous condensed phase and settled at the bottom of the tube. The dilute phase was subsequently decanted and replaced with more of the phase-separated mixture and allowed to undergo another cooling and droplet fusion cycle. This process was repeated until a sufficient condensed phase volume had formed that occupies the entirety of the NMR receiver coil with the dilute phase above, as shown in Fig. 3A. The phase-separated sample was allowed to equilibrate to the measurement temperature (25 or 40 °C) for ~8 hours prior to recording NMR experiments. After completion of condensed phase NMR measurements at each temperature, some of the dilute phase sitting above was decanted into a separate 3-mm NMR tube to generate the dilute phase samples used for measurements reported in Fig. 3. The labeling of the CAPRIN1 and FUS RRM components of the FUS RRM:CAPRIN1 condensate varied depending on the NMR experiment. The spectra shown in Fig. 3E and Fig. 4 (*blue*), for example, were measured using a condensate comprising $^{2}H$, $^{15}N$, $^{13}C$-ILV FUS RRM and $^{14}N$ CAPRIN1 ($^{14}N$ at 99.7% abundance), while the samples prepared for the experiments shown in Fig. 4 (*red*) and Fig. 5 consisted of $^{2}H$, $^{15}N$ FUS RRM and $^{14}N$ CAPRIN1 ($^{14}N$ at 99.99% abundance) where one-third of the CAPRIN1 molecules were also uniformly $^{13}C$-labeled. Another phase-separated sample consisting of $^{15}N$, $^{13}C$ FUS RRM and $^{14}N$ CAPRIN1 ($^{14}N$ at 99.7% abundance) was prepared for assignment of FUS RRM resonances in the condensed phase and for monitoring its aggregation (Fig. 3I).

**Circular dichroism (CD) thermal melts of RRMs**

CD thermal melts were performed on a Jasco J-1500 CD spectrometer using a 1 mm pathlength quartz cuvette. The CD signal at a wavelength of 218 nm was monitored over a temperature range of 30 °C to 80 °C, ramped at 1°C per minute. RRMs were prepared at 50 μM concentration either in 20 mM MES pH 6, 0.5 mM EDTA or 20 mM MES pH 5, 200 mM NaCl, 0.5 mM EDTA buffers. Spectra were smoothed with a first order Savitsky–Golay smoothing algorithm using a smoothing window of seven data points.

**NMR measurements**

NMR spectra were acquired on a 14.0-Tesla (600-MHz $^{1}H$ frequency) Bruker Avance III HD spectrometer, an 18.8 Tesla (800-MHz $^{1}H$ frequency) Bruker Avance III HD spectrometer, and a 23.5 Tesla (1 GHz $^{1}H$ frequency) Bruker Avance Neo spectrometer, all equipped with cryogenically cooled x, y, z pulsed-field gradient triple-resonance probes. An 11.7 Tesla (500-MHz $^{1}H$ frequency) Bruker Avance III HD spectrometer equipped with a liquid nitrogen-cooled *z* pulsed-field gradient triple-resonance probe was also used for monitoring RRM aggregation kinetics. Spectra were processed using NMRPipe(43) and analyzed with either NMRPipe, peakipy (https://github.com/j-brady/peakipy) or NMRFAM-SPARKY(44).

*Assignment of FUS RRM folded and unfolded backbone amide resonances*

Folded and unfolded FUS RRM backbone amide resonances were assigned using a suite of triple resonance 3D experiments, including 3D HNCO, HNCACO, HNCACB, HBCBCACONNH and HNN(45, 46), recorded using non-uniform sampling (NUS)(47) and processed with SMILE(48). Spectra were recorded at 40 °C on a mixed solution sample comprised of 0.9 mM $^{15}N$,$^{13}C$ FUS RRM and 9 mM CAPRIN1 in 20 mM MES pH 6, 0.5 mM EDTA, 10% $D_2O$(5, 6). Under the conditions of these experiments approximately 30% of FUS RRM is unfolded.

Assignments were transferred to the condensed phase by recording 3D HNCO and HNCACO experiments after phase-separating the sample with the addition of 200 mM NaCl, using the protocol described above.

*1D $^{15}$N-edited $^{1}$H NMR spectra for monitoring aggregation of RRMs (Fig. 1C-E)*

A timeseries of one dimensional $^{15}$N-edited $^{1}$H NMR spectra were recorded to monitor the aggregation kinetics of 0.5 mM $^{15}$N-labeled FUS, TAF-15 and EWSR1 RRMs in the absence and presence of 4 mM CAPRIN1. A 'dummy' sample containing a lower concentration of the protein in buffer was first used to optimize acquisition parameters so that the acquisition could begin shortly after insertion of the 'real' sample into the magnet. Spectra were recorded under fully relaxed conditions, *i.e.,* using a recycle delay of 5 s, and either 160 or 320 scans for total acquisition times of 15 or 30 mins, as needed to sufficiently sample the aggregation kinetics. The integral over the entire amide envelope was measured to determine the amount of NMR-visible RRM. Note that the concentration of natural abundance $^{15}$N-CAPRIN1 in the 4 mM CAPRIN1 samples is 12 μM so that the contribution from it to the measured integral can be neglected.

*Concentration-dependent 3D HNCO peak intensity measurements for determining intermolecular FUS RRM interaction sites (Fig. 2)*

Three-dimensional HNCO spectra were recorded at 45 °C on uniformly $^{15}$N, $^{13}$C-labeled FUS RRM prepared at 0.15 mM and 0.6 mM concentrations in 20 mM MES pH 6, 0.5 mM EDTA, 5% $D_2O$ buffer. Spectra were recorded with two scans, a recycle delay of 1.5 s, 17% NUS, and $^{15}$N and $^{13}$C acquisition times of 50 and 35 ms, respectively, for net experimental measurement times of ~5 hours. Peak intensities in the two 3D HNCO datasets were determined using NMRFAM-SPARKY. Intermolecular interactions were quantified by monitoring the decrease in HNCO peak intensities in the high ($I_{concentrated}$) *vs.* low ($I_{dilute}$) concentration datasets, after adjusting for the 4-fold concentration difference between samples.

*Measurement of water content in the dilute and condensed phases of phase-separated samples (Fig. 3B,D)*

The concentration of water, both $H_2O$ and $D_2O$, in the condensed and dilute phases was determined based on the integrals of the $H_2O$ ($D_2O$) peak in $^{1}$H ($^{2}$H) NMR spectra. Single-scan $^{1}$H NMR spectra were recorded at 25 °C on an 800 MHz spectrometer using a one-pulse acquire scheme with a low receiver gain to prevent receiver overflow, and an acquisition time of 64 ms. $^{1}$H pulses were centered on the water line, and a series of spectra were recorded with flip angles of {45°, 22.5°, 10°, 5°, 2.5°} and subsequently quantified. Similarly, single-scan $^{2}$H NMR spectra were recorded at 25 °C on a 500 MHz spectrometer using an acquisition time of 100 ms with $^{2}$H pulses centered on the water line. Comparison of $H_2O$ and $D_2O$ peak integrals in either the dilute or condensed phase samples with that of a reference buffer sample containing 90% $H_2O$ and 10% $D_2O$ was used to calculate the molar concentrations of water in each phase, as follows:

$$\left(\frac{V_{H2O,x}}{V_{H2O,Ref}}\right) x\ 0.9\ x\ 55\ \mathrm{M} + \left(\frac{V_{D2O,x}}{V_{D2O,Ref}}\right) x\ 0.1\ x\ 55\ \mathrm{M} \quad \text{[S1]}$$

where $V_{H2O,x}$ ($V_{D2O,x}$ ) is the integral of the $H_2O$ ($D_2O$) peak in either the dilute or condensed phase and $V_{H2O,Ref}$ ($V_{D2O,Ref}$) is the integral of the $H_2O$ ($D_2O$) peak in the reference sample. Note that concentrations of water calculated in this manner varied by less than 2 % as a function of $^{1}$H pulse flip angle.

*Measurement of the CAPRIN1 concentration in the FUS RRM:CAPRIN1 condensed phase via $^{1}$H NMR (Fig. 3C,D)*

The concentration of CAPRIN1 in the FUS RRM:CAPRIN1 condensed phase was determined by comparing peak integrals with a reference CAPRIN1 condensed phase sample of known concentration that lacks FUS RRM, as shown in Fig. 3C. Spectral regions from 1-2.1 ppm containing only proton peaks from CAPRIN1 were compared. Note that FUS RRM is deuterated with the exception of ILV methyl groups and backbone and sidechain amides; protons from methyls and amides resonate outside of the integrated region. Excitation-sculpting $^1$H NMR spectra (49) were recorded at 25 °C on an 800 MHz spectrometer with an interscan delay of 5 s, 128 scans, and spectral width and acquisition times of 16 ppm and 64 ms, respectively. The condensed phase concentration of CAPRIN1 in the reference sample was determined through $A_{280}$ measurements using an extinction coefficient of 10,430 $M^{-1}cm^{-1}$. A 2 µL aliquot extracted directly from the condensed phase was diluted 100-fold with 8M guanidinium chloride to generate a well-dispersed solution amenable to absorbance measurements.

*[$^1$H, $^{13}$C] methyl-TROSY ddHMQC NMR measurements of FUS RRM folded and unfolded populations and concentrations in CAPRIN1 dilute and condensed phases (Fig. 3F,G)*

Methyl-TROSY spectra were recorded of dilute and condensed phase samples at 25 and 40°C, 800 MHz, using a [$^1$H, $^{13}$C] ddHMQC pulse scheme with gradients for coherence transfer selection(21). Spectra were acquired under fully relaxed conditions with an interscan delay of 10 s, 8 scans/FID, $^{13}$C spectral widths, and acquisition times of 17 ppm and 25 ms, respectively, and a total acquisition time of 4 hours/spectrum. $^1$H and $^{13}$C pulses were centered on the water line and 19 ppm, respectively. Water suppression was obtained using coherence transfer selection gradients, with a water-selective EBURP1 pulse(50) (~ 7 ms) applied at the beginning of the scheme to ensure that the water signal is along the +Z axis at the start of $^1$H acquisition.

Integrals of the eight dispersed folded isoleucine resonances and the overlapping unfolded cluster were measured and the fractional populations of the folded and unfolded states, $P_F$ and $P_U$, respectively, were determined from the sum of the integrals of the folded and unfolded isoleucine resonances, accounting for differences in transverse relaxation of magnetization derived from the folded *vs.* unfolded species, as described previously(14):

$$P_F = \frac{\Sigma \frac{V_i^F}{exp\left(-R_{2,eff,i}^F * t_{seq}\right)}}{\Sigma \frac{V_i^F}{exp\left(-R_{2,eff,i}^F * t_{seq}\right)} + \Sigma \frac{V_T^U}{exp\left(-R_{2,eff}^U * t_{seq}\right)}} \quad \text{[S2]}$$

In Eq. [S2] $V_i^F$ is the volume of "folded" (F) peak *i*, and $V_T^U$ is the box-sum volume of the "unfolded" (U) peaks that cluster together, $t_{seq}$ is the length of all fixed delays during the [$^1$H,$^{13}$C] ddHMQC experiment (4.6 ms), $R_{2,eff,i}^F$ is an effective $^1$H single-quantum transverse relaxation rate for folded peak *i*, and $R_{2,eff}^U$ is the corresponding relaxation rate for the unfolded isoleucine cluster. Further details can be found in the *SI Appendix* of reference (14). The concentrations of the folded and unfolded FUS RRM species in the dilute and condensed phases were determined by comparing peak volumes with a FUS RRM sample in buffer of known concentration.

*1D $^{15}$N($^{13}$C)-edited $^1$H NMR spectra for monitoring aggregation of FUS RRM in buffer vs. dilute and condensed phases of FUS RRM:CAPRIN1 (Fig. 3I)*

Condensed and dilute phase samples of FUS RRM:CAPRIN1 were prepared with $^{15}$N,$^{13}$C FUS RRM and $^{14}$N CAPRIN1 ($^{14}$N at 99.7% abundance), using the protocol described above. Since the concentration of natural abundance $^{15}$N CAPRIN1 in the condensed phase is ~90 µM, of the same order as the FUS RRM concentration (488 µM at 25 °C), 1D $^{15}$N,$^{13}$C-edited (HNCO-based) $^1$H NMR spectra were recorded to selectively observe signals derived from FUS RRM. Integrals of the amide proton envelope in $^{15}$N,$^{13}$C-edited $^1$H NMR spectra recorded at 0 days *vs.* 89 days at 25 °C were compared to monitor the conversion of NMR visible FUS RRM protomers into NMR invisible aggregates. Measurements on the dilute phase sample were based on 1D $^{15}$N-edited $^{1}H$ NMR spectra (*i.e.,* without $^{13}C$ editing) since the concentration of natural abundance $^{15}N$ CAPRIN1 in the dilute phase is only ~8 μM and, therefore, signals from CAPRIN1 contribute negligibly to the measured integral. FUS aggregation kinetics in buffer were monitored using a 0.5 mM $^{15}N$,$^{13}C$ FUS RRM sample, approximately matching the condensed phase concentration of FUS RRM. As with the dilute phase, 1D $^{15}N$-edited $^{1}H$ NMR spectra were recorded at t=0 days and t=68 days for comparison.

*2D $^{1}H$-$^{15}N$ HSQC-based NOE experiment for mapping intermolecular FUS RRM:CAPRIN1 interactions in the condensed phase (Fig. 5)*

A condensed phase sample of FUS RRM:CAPRIN1 was prepared with $^{2}H$, $^{15}N$ FUS RRM and $^{14}N$ CAPRIN1 ($^{14}N$ at 99.99% abundance) where one-third of the CAPRIN1 molecules were also uniformly $^{13}C$-labeled. Using the pulse scheme outlined in *SI Appendix, Fig. S2* and described previously(22), NOEs were recorded in the condensed phase to measure intermolecular contacts between protons coupled to $^{13}C$ in the CAPRIN1 scaffold and protons coupled to $^{15}N$ in FUS RRM client molecules. NOESY datasets were recorded using a gradient selected TROSY readout(51) with an interscan delay of 1.5 s, 128 scans/FID, NOE mixing time of 250 ms, and $^{15}N$ $t_{1,max}$ = 50 ms, for a total acquisition time of ~20 hours per experiment. A control experiment recorded on a $^{2}H$, $^{15}N$ FUS RRM sample in buffer established that intra-molecular NOEs are not observed (*SI Appendix, Fig. S3*).

**Table S1 – Thermodynamics of the folding – unfolding equilibrium of FUS RRM in dilute *vs.* condensed phases**

| | **25 °C** | | | | | | | | **40 °C** | | | | | | | |
|---|---|---|---|---|---|---|---|---|---|---|---|---|---|---|---|---|
| **Phase** | $P_F$ % | $P_U$ % | **[F] μM** | **[U] μM** | **[T] μM** | $K_P$ | **$\Delta G_{F\to U}$ kJ/mol** | **$\Delta\Delta G_{F\to U}$ kJ/mol** | $P_F$ % | $P_U$ % | **[F] μM** | **[U] μM** | **[T] μM** | $K_P$ | **$\Delta G_{F\to U}$ kJ/mol** | **$\Delta\Delta G_{F\to U}$ kJ/mol** |
| **Dilute** | 92.3 | 7.7 | 201 | 17 | 218 | 2.2 | 6.2 | -4.2 | 74.6 | 25.4 | 155 | 53 | 208 | 2.5 | 2.8 | -4.3 |
| **Condensed** | 69.2 | 30.8 | 338 | 150 | 488 | | 2.0 | | 36.4 | 63.6 | 192 | 335 | 527 | | -1.5 | |

$$\Delta\Delta G_{F\to U}=\Delta G^{cond}_{F\to U} - \Delta G^{dil}_{F\to U}$$

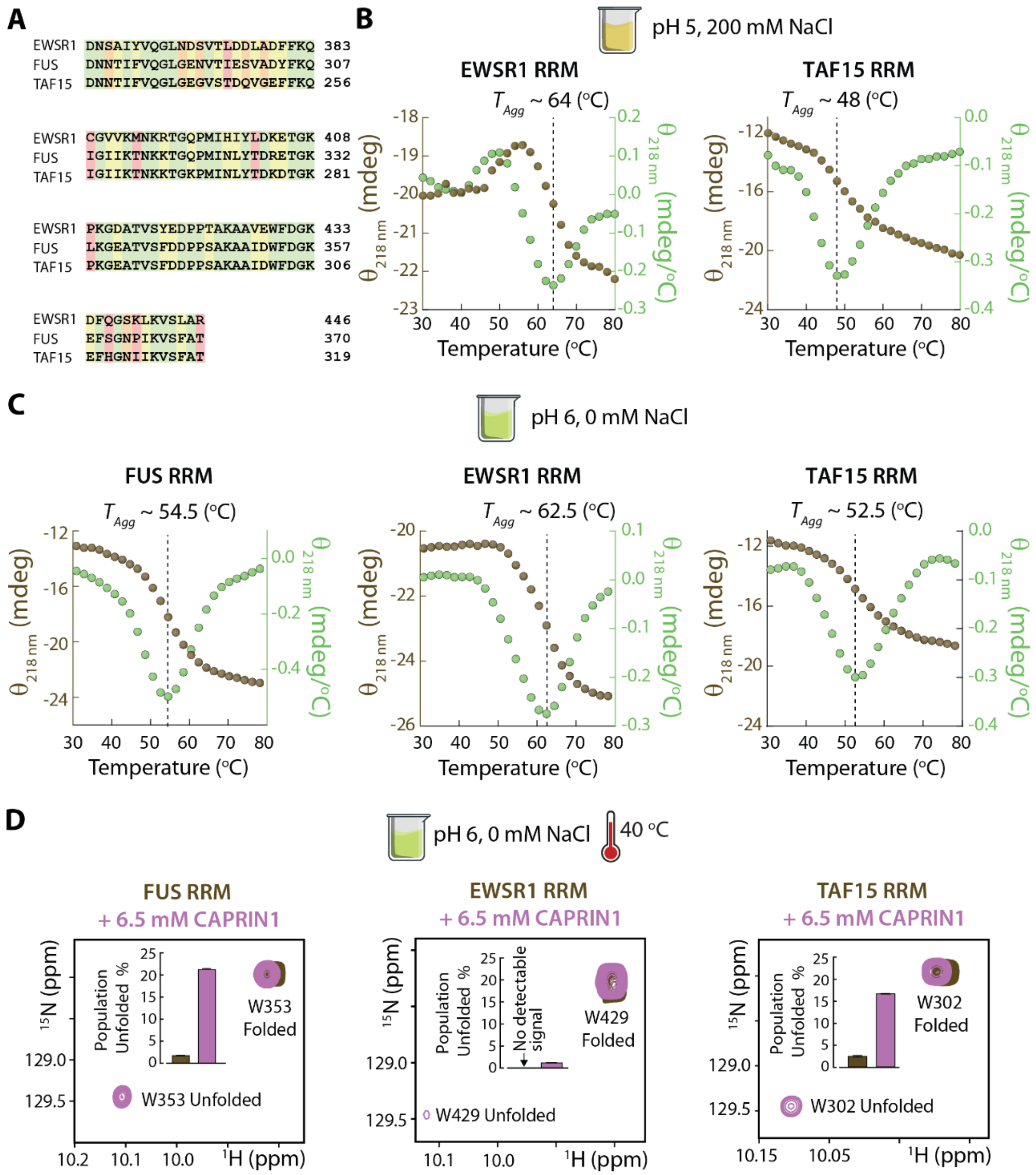


**Figure S1 – Thermal melts and CAPRIN1-induced unfolding of FUS, EWSR1, and TAF15 RRMs.** (**A**) Sequence alignments of FUS, EWSR1, and TAF15 RRMs, with green highlighting identical residues in all sequences, yellow signifying amino acids with similar properties at a given position, orange denoting weaker similarity between residues at a given position, and red indicating no significant similarities between residues. (**B**) Circular dichroism (CD) thermal melts monitoring the β-sheet signature at 218 nm (*brown*) of EWSR1 RRM (*left*) and TAF15 RRM (*right*) in MES pH 5 buffer with 200 mM NaCl. The derivatives of the thermal melt profiles are shown in green. (**C**) As (**B**), except for FUS RRM (*left*), EWSR1 RRM (*center*), and TAF15 RRM (*right*) in MES pH 6 buffer with 0 mM NaCl. (**D**) Zoomed-in expansions of $^{1}H$, $^{15}N$ TROSY-HSQC spectra of FUS (*left*), EWSR1 (*center*), and TAF15 (*right*) RRMs in the absence (*brown*) and presence (*purple*) of 6.5 mM CAPRIN1 highlighting the tryptophan indole peaks derived from the folded and unfolded states of the RRMs. Panel insets indicate the populations of the unfolded state, as determined from the peak volumes.

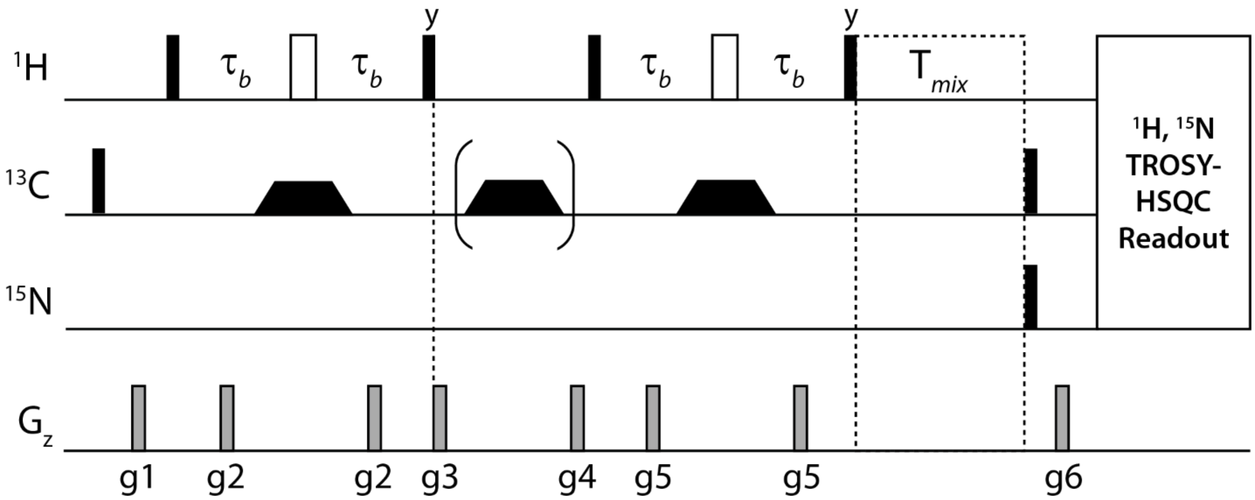


**Figure S2 – 2D $^1$H-$^{15}$N based NOE pulse scheme for recording intermolecular NOEs connecting $^{13}$C-bound protons of the CAPRIN1 scaffold and $^{15}$N-bound protons of the FUS client** (see above for labeling scheme). All rectangular $^1$H and $^{13}$C pulses are recorded at the highest possible power level, centered at the water frequency and at 67.5 ppm, respectively, while the $^{13}$C trapezoidal-shaped adiabatic pulses(52) have broadband excitation profiles covering aliphatic and aromatic regions of the carbon spectrum (400 μs, 80 kHz sweep centered at 67.5 ppm, 11 kHz maximum $B_1$ field). The second adiabatic pulse (in parenthesis) is applied in alternate scans with concomitant inversion of the receiver phase. The value of $\tau_b$ is set to 1.7 ms, while a mixing time, T*mix*, of 250 ms was used in the present set of experiments. The pulse sequence and parameters will be provided on Zenodo.

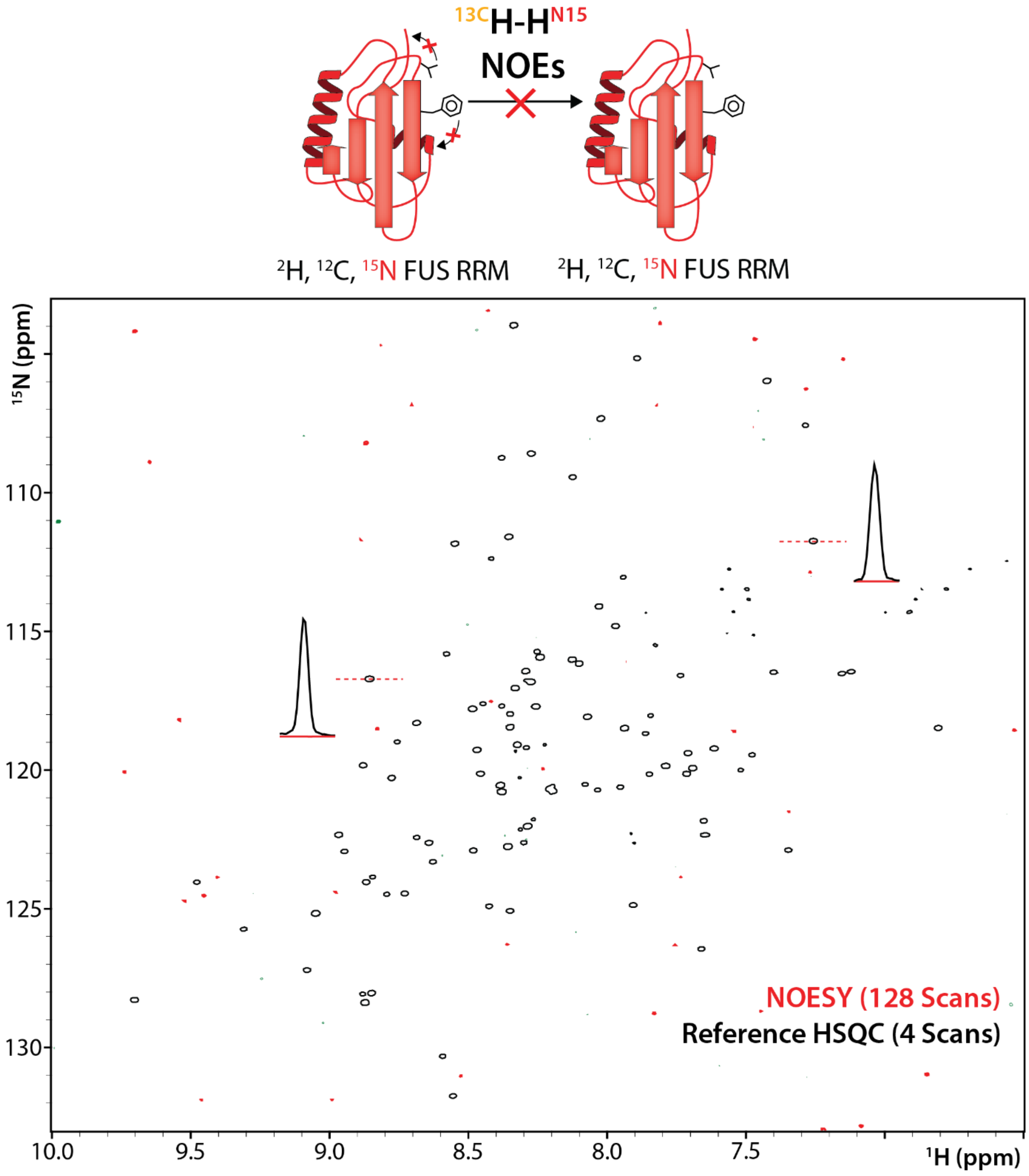


**Figure S3 – NOESY control experiment.** NOESY dataset (*red*) recorded using the scheme of Figure 2 with a mixing time of 250 ms at 1 GHz, 40 °C, is superimposed onto a reference $^{1}H$, $^{15}N$ TROSY-HSQC spectrum (*black, single contours*). The NOESY experiment is recorded with 32-fold more scans than the HSQC. Several traces are shown, centered at positions of peaks in the HSQC spectrum (black). The absence of corresponding NOESY peaks indicates that intramolecular NOE correlations are not observed using the labeling scheme described above. Thus, the peaks in the NOESY spectrum of Figure 5b of the main text are not artifactual and derive from magnetization transfer from CAPRIN1 to FUS RRM.
